# Getting personal: how vaccination exemptions shape herd immunity

**DOI:** 10.1101/500553

**Authors:** Emma R. Nedell, Romain Garnier, Saad B. Omer, Shweta Bansal

## Abstract

**Background:** State-mandated school entry immunization requirements in the United States play an important role in achieving high vaccine coverage and preventing outbreaks of vaccine-preventable diseases. Most states allow non-medical exemptions that let children remain unvaccinated on the basis of personal beliefs. However, the ease of obtaining such exemptions varies, resulting in a patchwork of state vaccination exemption laws, contributing to heterogeneity in vaccine coverage across the country. In this study, we evaluate epidemiological effects and spatial variations in non-medical exemption rates in the context of vaccine policies.

**Methods and Findings:** We first analyzed the correlation between non-medical exemption rates and vaccine coverage for three significant childhood vaccinations and found that higher rates of non-medical exemptions were associated with lower vaccination rates of school-aged children in all cases. We then identified a subset of states where exemption policy has recently changed and found that the effects on statewide non-medical exemption rates varied widely. Focusing further on Vermont and California, we illustrated how the decrease in non-medical exemptions due to policy change was concurrent to an increase in medical exemptions (in CA) or religious exemptions (in VT). Finally, a spatial clustering analysis was performed for Connecticut, Illinois, and California, identifying clusters of high non-medical exemption rates in these states before and after a policy change occurred. The clustering analyses show that policy changes affect spatial distribution of non-medical exemptions within a state.

**Conclusions:** Our work suggests that vaccination policies have significant impacts on patterns of herd immunity. Our findings can be used to develop evidence-based vaccine legislation.

## Introduction

Immunization requirements for school-entry date back to 1922 and have played a key role in achieving high levels of vaccine coverage against communicable diseases in the United States [1]. This patchwork of childhood immune protection, however, is punctured by a heterogeneous set of state-specific vaccination exemption rules. Medical exemptions to mandated vaccinations are available in all 50 states, and 47 offer non-medical exemptions in some form. Namely, 18 states offer personal belief exemptions for those who object to vaccinations for philosophical or moral reasons. In the remaining states offering non-medical exemptions, they are limited to religious beliefs. In the remaining three states (California, Mississippi, and West Virginia), only medical exemptions are available. While this has been the policy in Mississippi and West Virginia for decades [2], the ban on non-medical exemptions in California (enacted by CA Senate Bill 277 in January 2016) was motivated by the 2015 measles outbreak in the state [3] in which suboptimal vaccination rates in school-aged children was an important factor in the magnitude of the outbreak [4].

The ease of obtaining non-medical exemptions varies widely depending on state public health policies, from requiring a simple signature from the parents to obtaining a notarized document [5]. Generally, higher rates of non-medical exemptions are found in states where policies are more permissive [1, 6, 7]. In addition, states that allowed only religious exemptions, rather than religious and personal beliefs, have low non-medical exemption rates [8], although they tend to increase faster over time [6]. Policy efforts to slow down non-medical exemptions may also sometimes have unpredictable results. For instance, adopting a standardized form for exemption requests in order to better track exemptions may result in an increase in non-medical exemption rates [8]. This is because such a change may allow the emergence of resources facilitating the filing of exemptions by parents, resulting in effects opposite to the intended result.

The downstream impact of vaccination exemption policy on vaccination rates is also important to consider. Childhood vaccination rates tend to be lower in states with more permissive exemption policies [9], and a recent analysis has shown that in states allowing personal belief exemptions, higher levels of exemptions were associated with lower levels of measles, mumps, and rubella (MMR) vaccination in children attending kindergarten in the school year 2016-2017 [10]. Change in vaccination mandates increasing the difficulty for parents to obtain non-medical exemptions have had positive impacts on vaccination rates, including in Washington in 2011 [11] and California from the 2012-2013 school year [12]. However, assessing the association between policy changes and non-medical exemption rates remains necessary in other states and policy contexts. The success of California in eliminating non-medical exemptions comes in contrast with several failed legislative attempts in other states [9], even though the legality of non-medical exemption bans similar to the one implemented in California is not in question [13]. This variation in the success of legislative actions in reducing non-medical exemption rates demands that we assess variations in rates over consecutive years, in different policy and epidemiological contexts.

In this study, we focus on the epidemiological effects and spatial variations in non-medical exemption rates, and place it in the context of public health policies. We first assess the association between state-level non-medical exemptions and vaccination rates for three common childhood diseases, all mandatory for school-aged children. Next, we focus on the state-level dynamics of non-medical exemption rates over several school years in a subset of states that have implemented recent vaccination policy changes. Finally, we examine how spatial heterogeneity in non-medical exemption rates responds to policy changes at the county scale using four instances where legislative action to reduce accessibility to non-medical exemptions has recently been implemented, making them either harder to get or unavailable. Our analysis highlights how weak vaccination policies result in high non-medical exemption and low vaccination rates, producing hotspots of susceptible school-aged children for a number of vaccine-preventable infections. We advocate for aggressive public health policy changes to prevent further erosion in herd immunity for childhood diseases.

## Material and methods

To assess the association between non-medical exemption rates and vaccination rates at the beginning of school years 2016-2017 and 2017-2018, we used data in kindergarten from 48 states and the District of Columbia [14, 15]. No vaccination data were available for Oklahoma in 2016-2017, and neither vaccination or non-medical exemption rates were reported for Wyoming in both years. Oklahoma was thus excluded from the analysis in 2016-2017, and Wyoming was excluded in both years. Pennsylvania was also excluded from the analysis of the diphtheria, tetanus, and acellular pertussis (DTaP) vaccine in 2016-2017 because coverage data was not reported for this year [14]. We used a regression approach to test the associations between the proportion of non-medical exemption and vaccination rates. Because we are testing associations between proportions, we opted for a beta regression approach [16]. This analysis was run in R version 3.5.0 [17].

We identified a subset of states in which public health policies regarding non-medical exemptions have recently changed. Six states have made it harder to obtain non-medical exemptions between 2012 and 2016 [5]: Alaska (2013), Oregon (2014), Illinois (2015), Connecticut (2015), Missouri (2015), and Michigan (2015). In addition, in 2016, Vermont disallowed philosophical exemptions to only allow religious exemptions [18]. Finally, the state of California has strengthened its school immunization policies twice in the past decade: non-medical exemptions were made harder to obtain in 2013, and in 2015 new non-medical exemptions were barred from the beginning of the 2016-2017 school year [19]. For these states, we compiled data on non-medical exemptions in kindergartens from the Centers for Disease Control and Prevention (CDC) online annual school report results between 2003-2004 and 2010-2011, and from published annual surveys from school year 2011-2012 to school year 2017-2018 [20, 21, 22, 14, 23, 24, 15]. Data were not reported consistently for Illinois and Missouri for the period of 2012-2013 to 2016-2017. Because this period included the policy change, we did not include these two states in our analysis of policy changes. In addition, less than 10% of enrolled students were sampled in 2010-2011 and 2011-2012 in Alaska and these two years were not included in the dataset. We used a linear regression on years prior to the policy change to forecast NME rates in the absence of that change. In Vermont, we fitted the regression starting from school year 2008-2009, because of the sudden increase in NMEs during the school year 2007-2008. We would expect an effective policy change to lead to the data diverging from the forecasted trend.

Finally, we collected data on non-medical exemptions from state health departments at the county level in three states, including four instances of policy changes. In California, we obtained data on non-medical exemptions in kindergarten covering the period 2013-2014 up to 2016-2017, including two policy changes, at the beginning of the 2014-2015 and 2016-2017 school years respectively. In Connecticut, we included data on non-medical exemptions in kindergarten for the school year prior and the school year following a policy change in 2015, including school years 2014-2015 and 2015-2016. Finally, in Illinois, we compiled data on nonmedical exemptions in all school-aged children for the two years surrounding a policy change in 2015. Because Illinois only reports data for separate vaccines, exemptions specific for the MMR vaccine were used in that state. To analyze the spatial heterogeneity at the county level, and how the heterogeneity varied following policy changes, we computed *Moran’s I* [25] for each state and year. We performed a spatial clustering analysis for each state before and after the change in policy using SatScan [26] with the Bernoulli model [27, 28], as this model is adapted to our situation where individuals have or do not have an exemption. This method detects clusters of counties with high exemption rates relative to the rest of counties in a state; the mean rate of “high” clusters thus varies between states and between clusters. Maps were created in Python 3.6.3 using the Plotly graphing library package [29].

All data used in the manuscript, and codes for the statistical analysis are available on Github at github.com/Rom1Garnier/NME.

## Results

Folowing the analysis of Olive et al. [10], we expected a negative association between nonmedical exemptions and school-aged children vaccination levels. We extended their analysis for 2016-2017 to all states (irrespective of the breadth of the non-medical exemptions they allow) and found that higher rates of non-medical exemptions are associated with lower vaccination rates for the MMR vaccine (Figure 1A; *beta regression*; *p* < 0.001). Further, similar significant negative associations were present with two other common childhood vaccines included in the immunization mandates, DTaP (Figure 1B; *beta regression*; p = 0.002) and varicella (Figure 1C; *beta regression, p* = 0.02). We also obtained similar results for school year 2017-2018, with NMEs and vaccination rates being negatively associated (1D-F). Full results for the beta regressions can be found in Supplementary Table 1.

**Figure 1:**
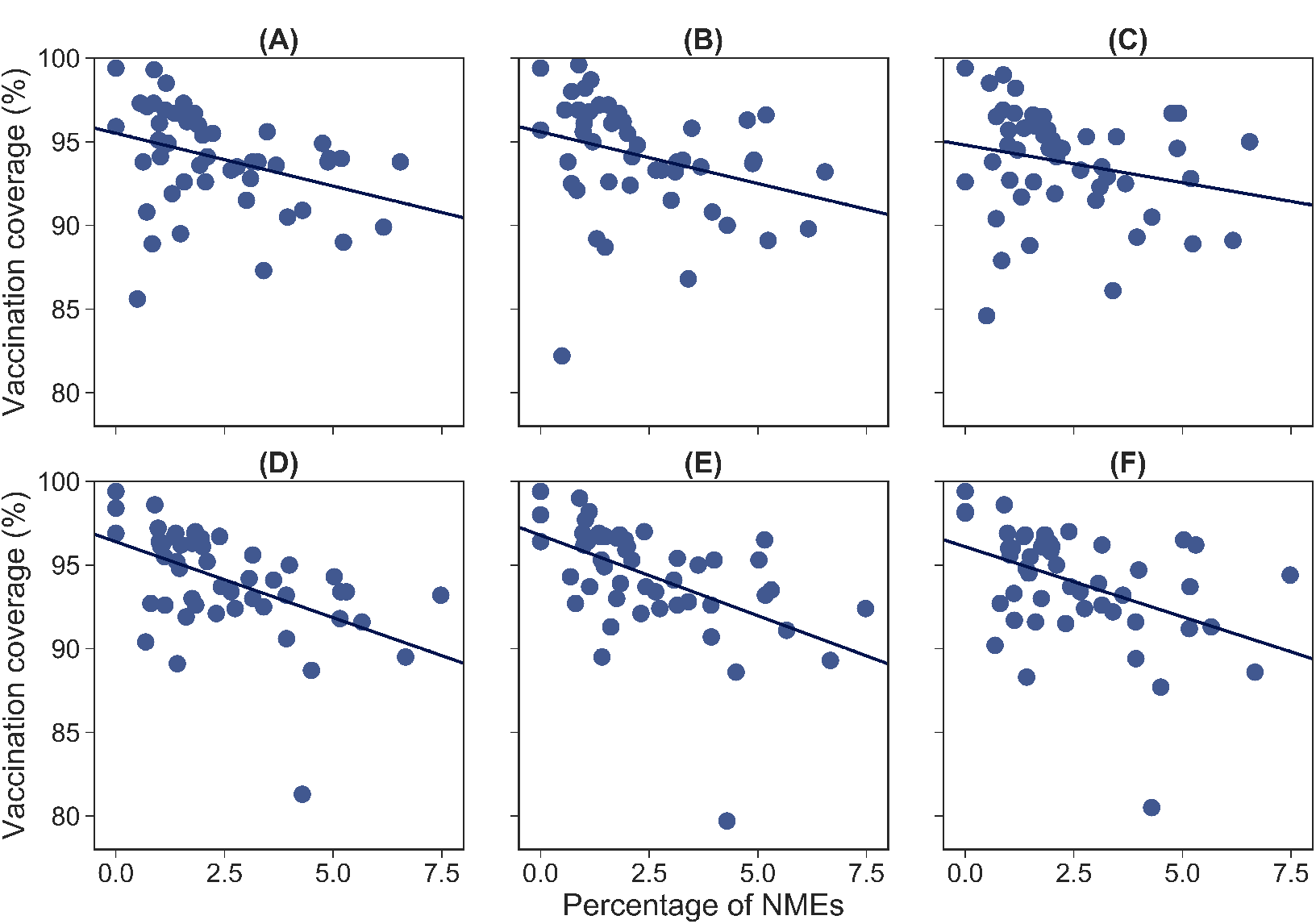
Association between percentages of non-medical exemptions and vaccination coverage at the state level in school year 2016-2017 (A-C) and school year 2017-2018 (D-F) for three common childhood vaccines: (**A**) Measles, Mumps, and Rubella (MMR); (**B**) Diphtheria, Tetanus, and acellular Pertussis (DTaP); (**C**) Varicella; (**D**) Measles, Mumps, and Rubella (MMR); (**E**) Diphtheria, Tetanus, and acellular Pertussis (DTaP); (**F**) Varicella.

We considered how changes in state public health policies affected non-medical exemption rates between school years 2011-2012 and 2017-2018, focusing on a set of six states which have implemented policy changes (Figure 2). First, we show that in one instance (Vermont, 2008), the levels of non-medical exemptions have increased rapidly from one year to the next. This was related to Vermont’s new requirement for immunization against hepatitis B and varicella (two doses) being enforced at the beginning of the 2008-2009 school year. Following this sudden increase, non-medical exemption rates showed no trend between 2008 and 2015 in Vermont (*linear regression, p* = 0.38). In all other states, non-medical exemption rates increased significantly from school year 2003-2004 until the considered policy change (*linear regression*, all *p* ≤ 0.007). The difference between the forecasted levels of non-medical exemptions and the actual non-medical exemption rates allows the identification of a number of situations (Figure 2A). Policies making it harder to obtain non-medical exemptions appear to have no apparent effect, with rates continuing to increase at apparently similar rates after the policy change (Alaska, Connecticut). In all the other cases, decreases were observed, with some being temporary (Oregon), and others seemingly more durable (California in 2014, Michigan). Finally, eliminating either the philosophical exemption in Vermont or non-medical exemptions altogether in California appears to have the strongest effect on the percentage of non-medical exemptions. However, in Vermont, the loss of philosophical belief exemptions was partly compensated by a sharp increase in religious exemptions, from 0.1% in school year 2014-2015 to 3.7% in 2016-2017 (Figure 2B). The decrease also appeared much slower in the second year after philosophical exemptions were banned. Similarly, in California, the sharp decrease in non-medical exemption rates was partly matched by a concurrent increase of medical exemptions from 0.17% in 2015-2016 to 0.51% in 2016-2017 (Figure 2B), probably in relation to how California Senate Bill 277 has provided for more physician discretion in the assessment of medical exemptions [19]. Reported exemption levels reached near zero as early as 2017-2018.

**Figure 2:**
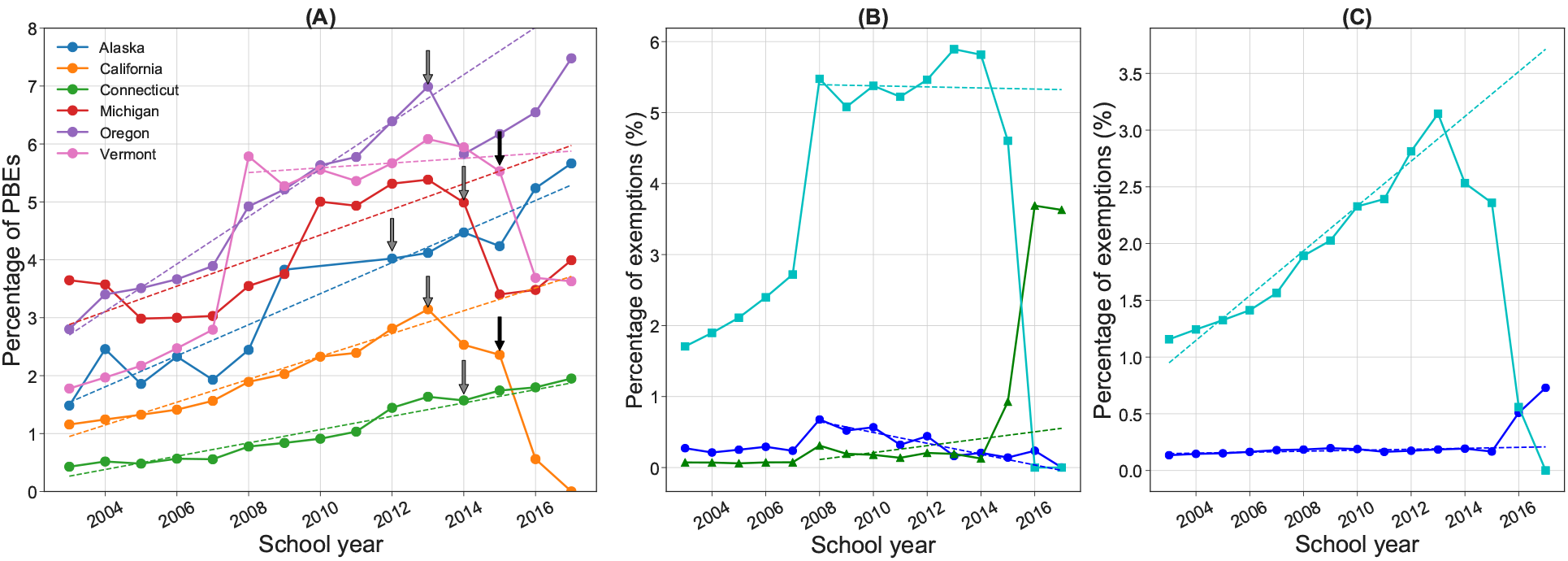
Dynamics of non-medical exemption rates between the school years 2003-2004 to 2016-2017. (**A**) In six states with recent exemption policy changes. The date of the policy change is indicated by an arrow. The solid line presents the data, while the dashed line represents the prediction of a linear regression fitted to the years prior to the first policy change in a state. The model was only fitted starting in 2008-2009 in Vermont. Black arrows indicate policies eliminating at least one type of NME; grey arrows indicate less stringent changes. (**B**) Dynamics of philosophical belief exemptions (light blue), religious exemptions (green), and medical exemptions (dark blue) in the state of Vermont. Solid lines represent the data, and dashed line represent predictions from a linear regression. (**C**) Details of the dynamics of total non-medical exemptions (light blue), and medical exemptions (dark blue) in the state of California. Solid lines represent the data, and dashed line represent predictions from a linear regression.

**Figure 3:**
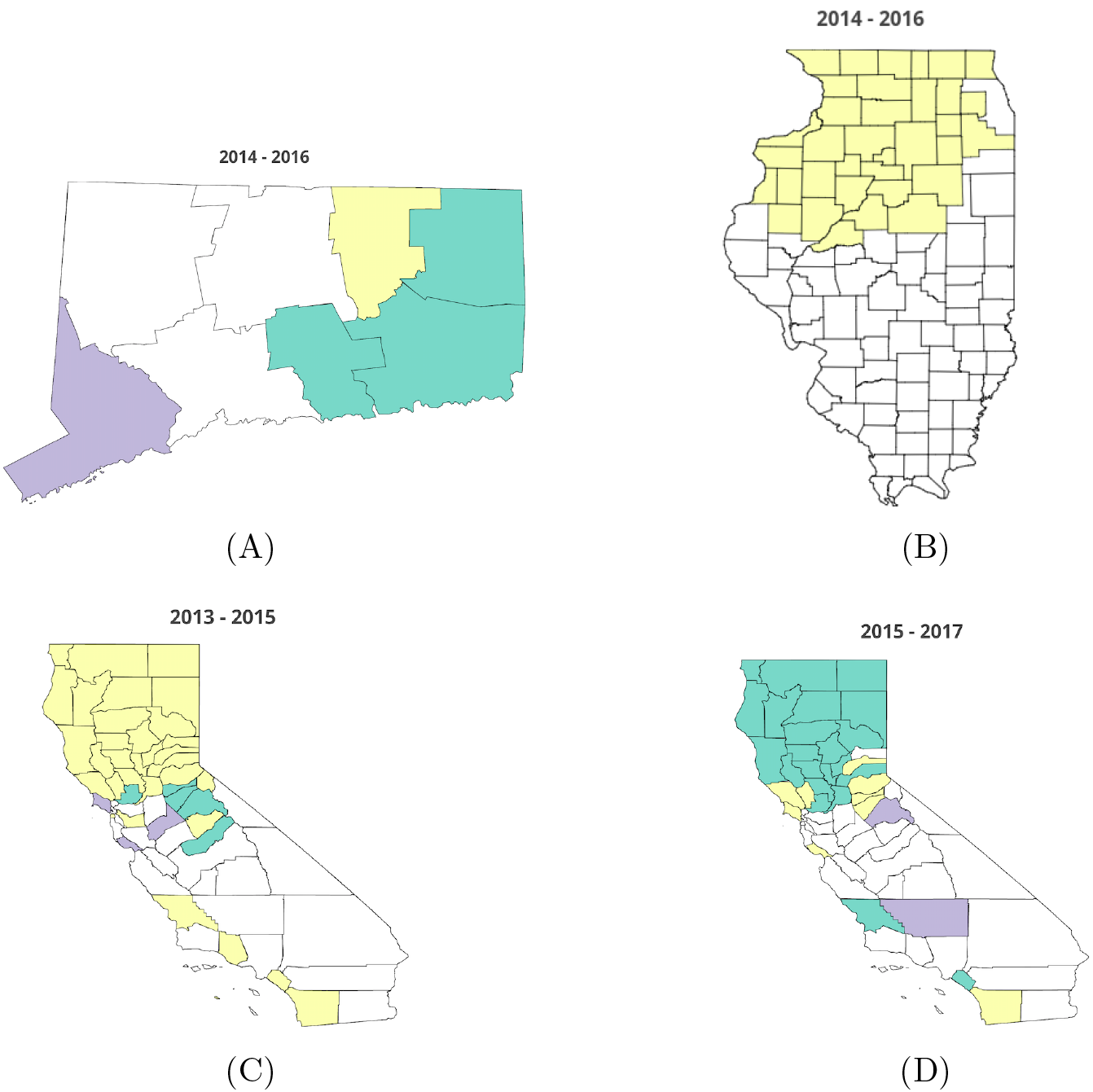
High non-medical exemption counties detected in a spatial clustering analysis performed for two school years surrounding a policy change. (**A**) Connecticut, school years 2014-2015 and 2015-2016; (**B**) Illinois, school years 2014-2015 and 2015-2016; (**C**) California, school years 2013-2014 and 2014-2015; (**D**) California, school years 2015-2016 and 2016-2017. Counties shaded in green belonged to a high non-medical exemption cluster only before the policy change; counties shaded in purple belonged to a high non-medical exemption cluster only after the policy change. Counties shaded in yellow belonged to a high non-medical exemption cluster both before and after the policy change.

Rates of non-medical exemptions in school-aged children showed spatial variability in all three states we focused on. However, we find that most policy changes have no significant effect on the mean and variance of non-medical exemption rates (Table 1). A reduction in both mean and variance of rates by county is only found between school years 2015-2016 and 2016-2017 in California, in relation to new non-medical exemptions becoming unavailable. We computed Moran’s I in all states and years (Table 1). In Illinois, we find that there is significant spatial heterogeneity in both years, with limited changes to Moran’s I before and after the policy change (school year 2014-2015: *Moran’s I:* 0.04; school year 2015-2016: *Moran’s I*: 0.05). In California, spatial heterogeneity remained significant before and after the first policy change in 2014 (i.e. making non-medical exemptions difficult but not eliminating them), which only resulted in a limited decrease of spatial heterogeneity indicated by a Moran’s I of 0.06 in school year 2013-2014 and a Moran’s I of 0.04 in 2014-2015. However, most significantly, the second policy change eliminating non-medical exemptions resulted in a loss of spatial heterogeneity. Indeed, we found significant spatial heterogeneity in school year 2015-2016 (*Moran’s I*: 0.08; *p <* 0.001) but Moran’s I becomes non-significant in school year 2016-2017 (*Moran’s I*: 0.01; *p* = 0.06). We find that there is no significant spatial heterogeneity in Connecticut both before and after the policy change. However, because Connecticut only has eight counties, this result needs to be taken with caution.

**Table 1:**
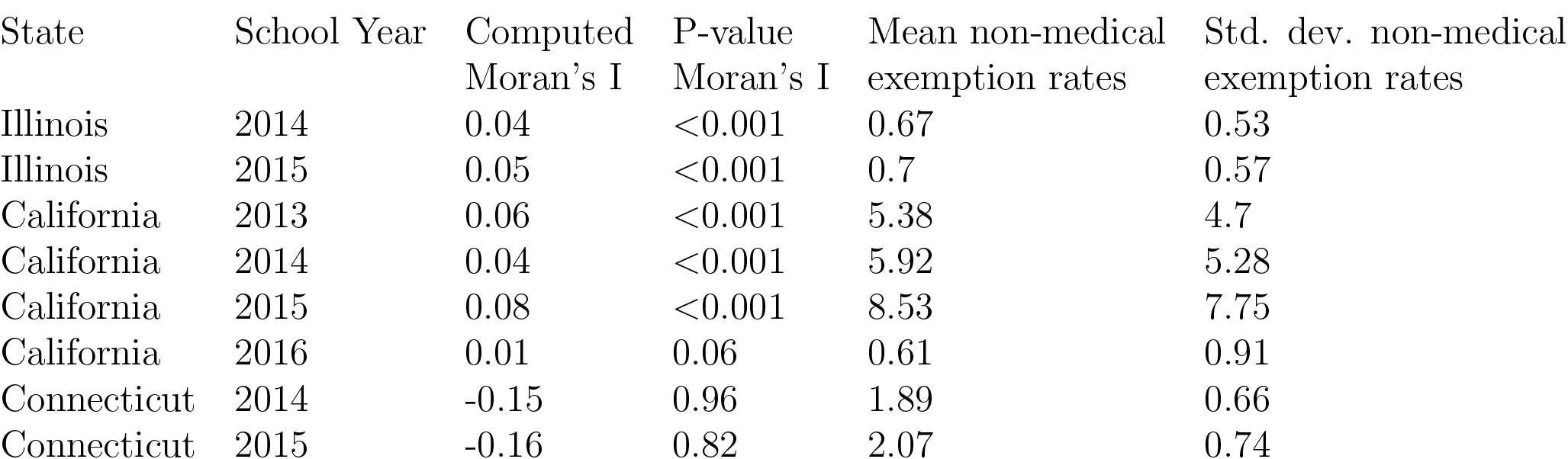
Computed Moran’s I, significance of Moran’s I, and mean and standard deviation of non-medical exemption rates for three states (California, Connecticut, Illinois) for which data was available at the county level. Policy changes occur in year 2015 in Illinois, year 2014 and 2016 in California, and in 2015 in Connecticut.

The spatial clustering analysis further shows how the policies impact the spatial distribution of non-medical exemptions (Figure 3, Supplementary Table 2). In Connecticut (Figure 3A), we identify different clusters between years indicating that spatial variation is present in both years, albeit with a shift in high risk groups. In Illinois (Figure 3B), the change in policy does not appear to have impacted the spatial clustering of non-medical exemptions. A single large cluster was identified in the northern part of the state both before and after the policy change.

Finally, in California, the two policy changes had different spatial impacts. The tightening of regulations around non-medical exemptions in 2014 appears to have had limited effects on spatial clustering of non-medical exemptions (Figure 3C), with a large cluster being identified in Northern California in both years. Conversely, this large cluster disappears in school year 2016-2017 and can only be identified in 2015-2016 (Figure 3D), indicating a large effect of Senate Bill 277, the legislation removing NMEs, on spatial heterogeneity in non-medical exemption rates. The large decrease in the mean percentage of exemptions of the remaining ‘high’ counties in California in 2016 further illustrates the effect of the policy change (Supplementary Table 2).

## Discussion

We have shown that, aggregated at the state level, non-medical exemption rates and vaccination rates are significantly associated for three major childhood vaccinations for which school immunization mandates exist. Furthermore, analyzing the dynamics of non-medical exemption in several states with policy change history, we showed that eliminating either a subset of exemptions (as in Vermont) or all non-medical exemptions (as in California) appears most effective in reducing exemption rates overall. Finally, we showed that non-medical exemptions are clustered at the county level, and that only the most stringent policy change appeared to modify both the spatial heterogeneity and the mean and variance in non-medical exemption rates.

The association between childhood vaccination rates and non-medical exemptions has important implications for vaccine-preventable childhood infectious disease risk. This association, along with the heterogeneous spatial distribution of non-medical exemptions, creates pockets of eroded herd immunity where outbreaks of vaccine preventable diseases would be more likely [30]. Furthermore, we illustrate that this is true not only for MMR [10] but for a wide range of childhood diseases. It is thus important to consider the compounded risk of all childhood diseases when evaluating the public health risk posed by non-medical exemptions. Individuals with non-medical exemptions have an increased risk of contracting vaccine-preventable diseases such as measles, and higher rates of exempted individuals in the population can increase the incidence of the disease in vaccine-protected populations [31]. Intentionally unvaccinated individuals indeed make up large proportions of cases in outbreaks of both measles and pertussis in the United States [32], and can unwittingly be the starting point of epidemics that may take hold in population with relatively high vaccination rates [33]. The potential co-circulation of childhood infections also raises concerns of immunological and ecological interference between the diseases [34, 35].

We highlight that policies that reduce the spatial heterogeneity and variance in non-medical exemption rates are key to eliminating pockets of susceptibility and minimizing the risk of childhood disease outbreaks. Our work suggests that making non-medical exemptions more difficult to obtain by increasing the administrative burden for parents is unlikely to achieve this goal. Only the complete removal of non-medical exemptions in California shows promise and may represent an effective policy tool. A similar spatial analysis of Vermont would be needed to assess whether the partial removal of NMEs has similar spatial effects. Additionally, we highlight that data at finer spatial scales could reveal the presence of these spatial effects below the county-scale.

We note that it is important to account for the effects of grand-fathered exemptions, i.e. in case of new laws restricting or eliminating exemptions, allowing children with existing exemptions to maintain their exempt status. Therefore, it may indeed take several years for existing non-medical exemptions to be grand-fathered, and, in the case of California, a zero non-medical exemption rate was estimated to only be achieved in 2022 even though no new non-medical exemptions have been granted since the beginning of school year 2016-2017 [36]. This means that return to optimal herd immunity levels may take several years.

However, the data available for the 2017-2018 school year indicates that NMEs are already at near zero, with only 5 NMEs left in the entire state [15]. The immediate benefits may also markedly differ depending on whether non-medical exemptions are granted for several years (as was the case in California) or whether they require annual renewal because of state or school policies [37].

We also argue that the context of what alternative exemptions are available to parents when access to some exemptions becomes more difficult needs to be taken into account to maximize the increase in vaccination coverage. Indeed, both the increase in religious exemptions in Vermont and in medical exemptions in California points towards parents seeking alternative exemptions whenever possible. The positive relationship between increase in medical exemptions and past rates of non-medical exemptions at the county level in California also supports this idea [19]. An increase in medical exemption could be expected in response to any increase in the difficult of obtaining non-medical exemptions [11]. However, states where non-medical exemptions are hard to obtain have only slightly higher medical exemption rates if the procedure to obtain these exemptions remains stringent [38]. The simplification of the medical exemption process in California, introduced in Senate Bill 277 alongside the elimination of non-medical exemptions, may thus be partly to blame for the sharp increase in medical exemptions at the start of the 2016-2017 school year [19, 39]. While the child’s healthcare professional is often in the best position to offer relevant counsel on immunization to vaccine-hesitant parents [40], parents may put pressure on providers to obtain medical exemptions and/or turn to more sympathetic providers [11]. Additionally, recent studies have shown a rise in conditional admissions after an exemption policy change [11] (which is not something we included in our analysis), thus further consideration of effect of this category of students is also needed [12]. Variable proportions of conditional admissions could, for instance, partly explain the noise in the association between NME rates and vaccination rates. We argue that, in order to maximize the effects of the elimination of (some) exemptions, efforts should be made to keep other types at least as difficult to obtain as they were prior to the new policy.

More generally, the question of whether a model with only medical exemptions would be well accepted and/or enforceable in the United States is an open question [2, 41]. Monetary incentives have been suggested to discourage parents from obtaining non-medical exemptions, in particular in the form of fees [42]. The rationale is that fees would reduce the convenience of non-medical exemptions and result in increase of vaccination rates, while any money collected would help alleviate the financial burden that vaccine-exempt individuals place on taxpayers. Another possible option, used for instance in Australia, could be to tie welfare payments to children vaccination records [43]. However, in the context of the United States, this policy could be misguided: vaccine refusal has been shown to be more prevalent in higher socio-economic neighborhoods [44] where welfare payments may be uncommon. From an ethical standpoint, which approach is preferable between making non-medical exemptions harder to get through administrative or time-consuming hurdles, and outright elimination of non-medical exemptions is far from settled [45, 46]. Even though there is a strong legal basis that would allow states to ban non-medical exemptions [13], partial elimination targeting diseases whose transmission is primarily school based such as measles may be preferable to avoid further strengthening anti-vaccine sentiments [40]. Communication around the benefits and safety of vaccines should represent a key component of any elimination effort, even though education of vaccine-refusing parents has proven challenging [47]. In any case, while the exploration of models used in other countries around the world provides useful data, understanding the local and national context is likely to be key to the implementation of a successful policy aimed at maximizing vaccination rates and herd immunity [48].

The benefits of herd immunity for childhood infections cannot be overstated. The reduction of non-medical exemption rates through NME policies remains a powerful tool in the fight to maintain herd immunity. However, effective policies regarding vaccination exemptions require careful evaluation of the relative costs and benefits in the near- and long-term.

## Acknowledgments

Research reported in this publication was supported by the National Institute Of General Medical Sciences of the National Institutes of Health under Award Number R01GM123007. The content is solely the responsibility of the authors and does not necessarily represent the official views of the National Institutes of Health.

## Supplementary-Material

**Supplementary Table 1:**
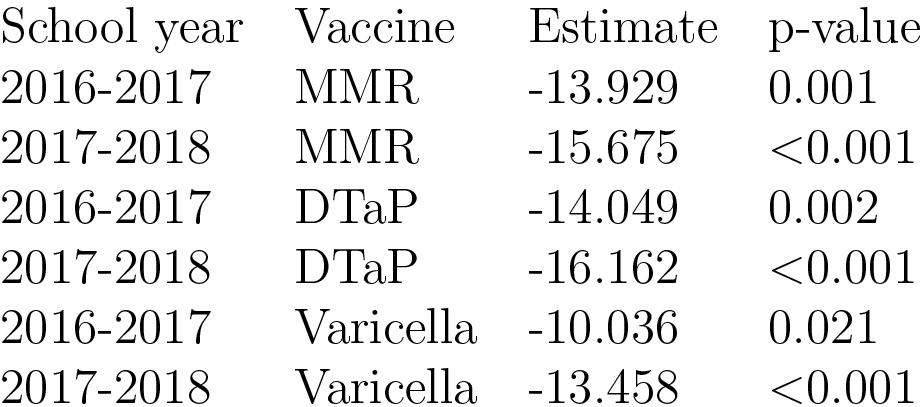
Estimates and p-values from the beta regression of NME rates and vaccination rates for each vaccine and school year at the state-level. All the associations are significant and negative, supporting a negative association between NMEs and vaccination rates.

**Supplementary Table 2:**
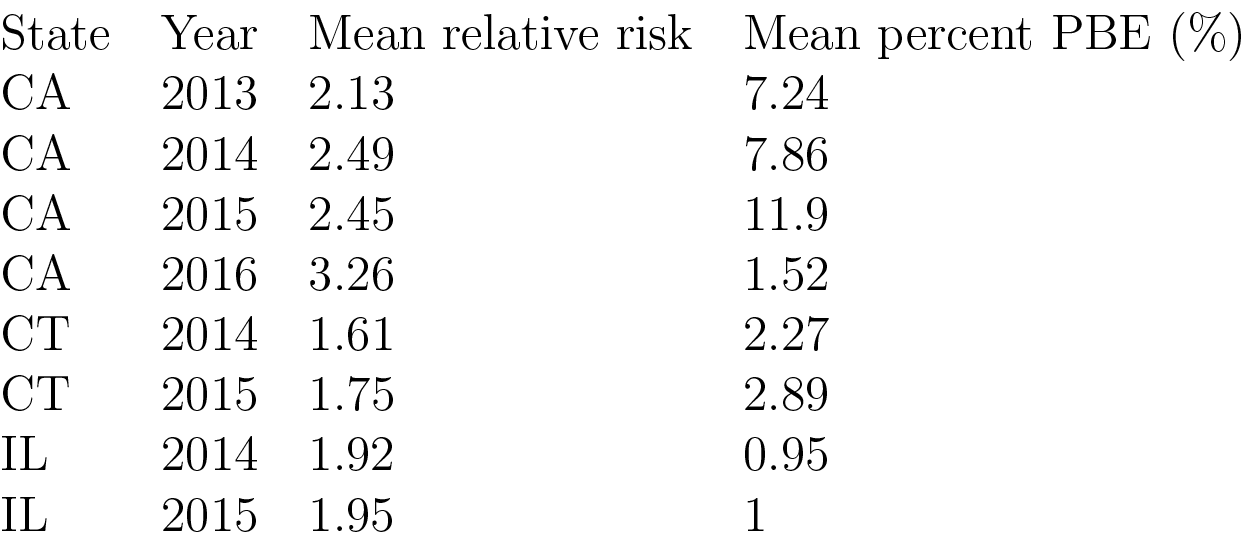
Mean relative risk and mean percentages of personal belief exemptions (PBE) in clusters of “high” PBE rates detected using SatScan. Mean relative risk corresponds to the average of risk of high PBEs in counties detected as “high” risk by the SatScan algorithm relative to the rest of the counties in a given state and year.

| State | Year | Mean relative risk | Mean percent PBE (%) |
| --- | --- | --- | --- |
| CA | 2013 | 2.13 | 7.24 |
| CA | 2014 | 2.49 | 7.86 |
| CA | 2015 | 2.45 | 11.9 |
| CA | 2016 | 3.26 | 1.52 |
| CT | 2014 | 1.61 | 2.27 |
| CT | 2015 | 1.75 | 2.89 |
| IL | 2014 | 1.92 | 0.95 |
| IL | 2015 | 1.95 | 1 |

## References

[1] E. Wang, J. Clymer, C. Davis-Hayes, and A. Buttenheim. Nonmedical exemptions from school immunization requirements: a systematic review. Am J Public Health, 104(11):e62–e84, 2014.

[2] J. Colgrove and A. Lowin. A tale of two states: Mississippi, West Virginia, and exemptions to compulsory school vaccination laws. Health Aff (Millwood), 35(2):348–55, 2016.

[3] J. Zipprich, K. Winter, J. Hacker, D. Xia, J. Watt, and K. Harriman. Measles outbreak-California, december 2014 - february 2015. MMWR Morb Mortal Wkly Rep, 64(6):153–154, 2015.

[4] M. S. Majumder, E. L. Cohn, S. R. Mekaru, J. E. Huston, and J. S. Brownstein. Substandard vaccination compliance and the 2015 measles outbreak. JAMA Pediatr, 169(5):494–5, 2015.

[5] S. B. Omer, R. M. Porter, K. Allen, D. A. Salmon, and R. A. Bednarczyk. Trends in kindergarten rates of vaccine exemption and state-level policy, 2011-2016. Open Forum Infect Dis, 5(2):ofx244, 2018.

[6] S. B. Omer, J. L. Richards, M. Ward, and R. A. Bednarczyk. Vaccination policies and rates of exemption from immunization, 2005-2011. N Engl J Med, 367(12):1170–1, 2012.

[7] S. B. Omer, W. K. Y. Pan, N. A. Halsey, S. Stokley, L. H. Moulton, A. M. Navar, M. Pierce, and D. A. Salmon. Nonmedical exemptions to school immunization requirements-secular trends and association of state policies with pertussis incidence. JAMA, 296(14):1757–1763, 2006.

[8] W. D. Bradford and A. Mandich. Some state vaccination laws contribute to greater exemption rates and disease outbreaks in the United States. Health Aff (Millwood), 34(8):1383–90, 2015.

[9] J. Shaw, E. M. Mader, B. E. Bennett, O. K. Vernyi-Kellogg, Y. T. Yang, and C. P. Morley. Immunization mandates, vaccination coverage, and exemption rates in the United States. Open Forum Infect Dis, 5(6):ofy130, 2018.

[10] J. K. Olive, P. J. Hotez, A. Damania, and M. S. Nolan. The state of the antivaccine movement in the United States: A focused examination of nonmedical exemptions in states and counties. PLoS Med, 15(6):e1002578, 2018.

[11] S. B. Omer, K. Allen, D. H. Chang, L. B. Guterman, R. A. Bednarczyk, A. Jordan, A. Buttenheim, M. Jones, C. Hannan, M. P. deHart, and D. A. Salmon. Exemptions from mandatory immunization after legally mandated parental counseling. Pediatrics, 141(1):e20172364, 2018.

[12] A. M. Buttenheim, M. Jones, C. McKown, D. Salmon, and S. B. Omer. Conditional admission, religious exemption type, and nonmedical vaccine exemptions in california before and after a state policy change. Vaccine, 36(26):3789–3793, 2018.

[13] M. M. Mello, D. M. Studdert, and W. E. Parmet. Shifting vaccination politics - the end of personal-belief exemptions in california. N Engl J Med, 373(9):785–787, 2015.

[14] R. Seither, K. Calhoun, E. J. Street, J. Mellerson, C. L. Knighton, A. Tippins, and J. M. Underwood. Vaccination coverage for selected vaccines, exemption rates, and provisional enrollment among children in kindergarten-United States, 2016-17 school year. MMWR Morb Mortal Wkly Rep, 66(40):1073–1080, 2017.

[15] J. L. Mellerson, C. B. Maxwell, C. L. Knighton, J. L. Kriss, R. Seither, and C. L. Black. Vaccination coverage for selected vaccines and exemption rates among children in kindergarten-united states 2017-2018 school year. Morbidity and Mortality Weekly Report (MMWR), 67(40):1115–1122, 2018.

[16] Francisco Cribari-Neto and Achim Zeileis. Beta regression in R. J Stat Softw, 34(2):1–24, 2010.

[17] R Core Team. R: A Language and Environment for Statistical Computing. R Foundation for Statistical Computing, Vienna, Austria, 2013.

[18] Vermont Department of Health. Vermont immunization program - 2017 annual report. Report, Vermont Department of Health, 2017.

[19] P. L. Delamater, T. F. Leslie, and Y. T. Yang. Change in medical exemption from immunization in california after elimination of personal belief exemptions. JAMA, 318(9):862–863, 2017.

[20] S. M. Greby, K. G. Wooten, C. L. Knighton, B. Avey, and S. Stokley. Vaccination coverage among children in kindergarten-United States, 2011-12 school year. MMWR Morb Mortal Wkly Rep, 61(33):647–652, 2012.

[21] R. Seither, K. Calhoun, C. L. Knighton, J. Mellerson, S. Meador, A. Tippins, S. M. Greby, and V. Dietz. Vaccination coverage among children in kindergarten-United States, 2014-15 school year. MMWR Morb Mortal Wkly Rep, 64(33):897–904, 2015.

[22] R. Seither, K. Calhoun, J. Mellerson, C. L. Knighton, E. Street, V. Dietz, and J. M. Underwood. Vaccination coverage among children in kindergarten-United States, 2015-16 school year. MMWR Morb Mortal Wkly Rep, 65(39):1057–1064, 2016.

[23] R. Seither, S. Masalovich, C. L. Knighton, J. Mellerson, J. A. Singleton, and S. M. Greby. Vaccination coverage among children in kindergarten-United States, 2013-14 school year. MMWR Morb Mortal Wkly Rep, 63(41):913–920, 2014.

[24] R. Seither, L. Shaw, C. L. Knighton, S. M. Greby, and S. Stokley. Vaccination coverage among children in kindergarten-United States, 2012-13 school year. MMWR Morb Mortal Wkly Rep, 62(30):607–612, 2013.

[25] E. Paradis, J. Claude, and K. Strimmer. APE: analyses of phylogenetics and evolution in R language. Bioinformatics, 20:289–290, 2004.

[26] M. Kulldorff. SaTScan v9.6: Software for the spatial and space-time scan statistics, 2018.

[27] C. Aloe, M. Kulldorff, and B. R. Bloom. Geospatial analysis of nonmedical vaccine exemptions and pertussis outbreaks in the United States. Proc Natl Acad Sci USA, 114(27):7101–7105, 2017.

[28] M. Kulldorff. A spatial scan statistic. Communications in Statistics: Theory and Methods, 26:1481–1496, 1997.

[29] Plotly Technologies Inc. Collaborative data science. Plotly Technologies Inc., Montreal, QC, 2015.

[30] S. B. Omer, D. A. Salmon, W. A. Orenstein, M. P. DeHart, and N. Halsey. Vaccine refusal, mandatory immunization, and the risks of vaccine-preventable diseases. N Engl J Med, 360:1981–8, 2009.

[31] D. A. Salmon, M. Haber, E. J. Gangarosa, L. Phillips, N. J. Smith, and R. T. Chen. Health consequences of religious and philosophical exemptions from immunization laws. individual and societal risks of measles. JAMA, 281(1):47–54, 1999.

[32] V. K. Phadke, R. A. Bednarczyk, D. A. Salmon, and S. B. Omer. Association between vaccine refusal and vaccine-preventable diseases in the United States: A review of measles and pertussis. JAMA, 315(11):1149–58, 2016.

[33] D. E. Sugerman, A. E. Barskey, M. G. Delea, I. R. Ortega-Sanchez, D. Bi, K. J. Ralston, P. A. Rota, K. Waters-Montijo, and C. W. Lebaron. Measles outbreak in a highly vaccinated population, san diego, 2008: role of the intentionally undervaccinated. Pediatrics, 125(4):747–55, 2010.

[34] Diane E Griffin. Measles virus-induced suppression of immune responses. Immunol Rev, 236(1):176–189, 2010.

[35] P Rohani, CJ Green, NB Mantilla-Beniers, and BT Grenfell. Ecological interference between fatal diseases. Nature, 422(6934):885, 2003.

[36] P. L. Delamater, T. F. Leslie, and Y. T. Yang. A spatiotemporal analysis of non-medical exemptions from vaccination: California schools before and after SB277. Soc Sci Med, 168:230–238, 2016.

[37] D. A. Salmon, S. B. Omer, L. H. Moulton, S. Stokley, M. P. DeHart, S. Lett, B. Norman, S. Teret, and N. A. Halsey. Exemptions to school immunization requirements: the role of school-level requirements, policies, and procedures. Am J Public Health, 95(3):436–440, 2005.

[38] S. Stadlin, R. A. Bednarczyk, and S. B. Omer. Medical exemptions to school immunization requirements in the United States-association of state policies with medical exemption rates (2004-2011). J Infect Dis, 206(7):989–92, 2012.

[39] S. Mohanty, A. M. Buttenheim, C. M. Joyce, A. C. Howa, D. Salmon, and S. B. Omer. Experiences with medical exemptions after a change in vaccine exemption policy in california. Pediatrics, 142(5):e20181051, 2018.

[40] D. J. Opel, J. L. Schwartz, S. B. Omer, R. Silverman, J. Duchin, E. Kodish, D. S. Diekema, E. K. Marcuse, and W. Orenstein. Achieving an optimal childhood vaccine policy. JAMA Pediatr, 171(9):893–896, 2017.

[41] D. J. Opel, M. P. Kronman, D. S. Diekema, E. K. Marcuse, J. S. Duchin, and E. Kodish. Childhood vaccine exemption policy: The case for a less restrictive alternative. Pediatrics, 137(4), 2016.

[42] J. K. Billington and S. B. Omer. Use of fees to discourage nonmedical exemptions to school immunization laws in U.S. states. Am J Public Health, 106(2):269–70, 2016.

[43] Y. T. Yang and D. M. Studdert. Linking immunization status and eligibility for welfare and benefits payments: the australian “no jab, no pay” legislation. JAMA, 317(8):803–804, 2017.

[44] Sandra Goldlust, Elizabeth C. Lee, Murali Haran, Pejman Rohani, and Shweta Bansal. Assessing the distribution and determinants of vaccine underutilization in the United States. BioRXiv, 2017.

[45] Mark Christopher Navin and Mark Aaron Largent. Improving nonmedical vaccine exemption policies: Three case studies. Public Health Ethics, 10(3):225–234, 2017.

[46] A. Giubilini, T. Douglas, and J. Savulescu. Liberty, fairness, and the ‘contribution model’ for non-medical vaccine exemption policies: a reply to navin and largent. Public Health Ethics, 10(3):235–240, 2017.

[47] M. C. Navin, A. T. Kozak, and E. C. Clark. The evolution of immunization waiver education in michigan: A qualitative study of vaccine educators. Vaccine, 36(13):1751–1756, 2018.

[48] K. Attwell, M. C. Navin, P. L. Lopalco, C. Jestin, S. Reiter, and S. B. Omer. Recent vaccine mandates in the united states, europe and australia: A comparative study. Vaccine, 36(48):7377–7384, 2018.

